# A comparison between low-cost library preparation kits for low coverage sequencing

**DOI:** 10.1101/2024.01.30.578044

**Authors:** Caitlin M. Stewart, Matthew JS Gibson, Jahan-Yar Parsa, Jeremiah H. Li

## Abstract

In the fields of human health and agricultural research, low coverage whole-genome sequencing followed by imputation to a large haplotype reference panel has emerged as a cost-effective alternative to genotyping arrays for assaying large numbers of samples. However, a systematic comparison of library preparation methods tailored for low coverage sequencing remains absent in the existing literature. In this study, we evaluated one full sized kit from IDT and miniaturized and evaluated three Illumina-compatible library preparation kits—the KAPA HyperPlus kit (Roche), the DNA Prep kit (Illumina), and an IDT kit—using 96 human DNA samples. Metrics evaluated included imputation concordance with high-depth genotypes, coverage, duplication rates, time for library preparation, and additional optimization requirements. Despite slightly elevated duplication rates in IDT kits, we find that all four kits perform well in terms of imputation accuracy, with IDT kits being only marginally less performant than Illumina and Roche kits. Laboratory handling of the kits was similar: thus, the choice of a kit will largely depend on (1) existing or planned infrastructure, such as liquid handling capabilities, (2) whether a specific characteristic is desired, such as the use of full-length adapters, shorter processing times, or (3) use case, for instance, long vs short read sequencing. Our findings offer a comprehensive resource for both commercial and research workflows of low-cost library preparation methods suitable for high-throughput low coverage whole genome sequencing.

## INTRODUCTION

Large scale genotyping studies have historically relied on array based methods due to the relative expense of sequencing (LaFramboise 2009). In comparison to arrays, recent work has shown that low coverage sequencing, on the order of 0.5-1X coverage, combined with imputation can be a cost effective and superior alternative (Chat *et al*. 2022, Li *et al*. 2021). In addition, low coverage whole genome sequencing can identify relevant single nucleotide polymorphisms (SNPs) not covered by arrays, increase the power of genome wide association studies (GWAS), be used in place of other methods for pharmacogenomic studies, and decrease measurement errors in polygenic risk scores (PRS) (Chat *et al*. 2022; Gilly *et al*. 2019; Homburger *et al*. 2019; Li *et al*. 2021; Wasik *et al*. 2021). For DNA sequencing, one of the most critical components is the choice of library preparation methods. While many studies exist for the selection of library preparation methods for high-coverage sequencing (Aigrain 2016; Ribarska 2022), researchers seeking to do low coverage whole genome sequencing frequently make decisions at this critical step without guidance. The absence of publicly available guidelines for low coverage library preparations exacerbates this issue.

Considerations in choosing a library preparation workflow include cost and time as well as data quality metrics including sequencing duplication rates, sequencing coverage, and, in the case of low coverage sequencing, imputation accuracy. We chose four methods to compare. These library preparation methods are from commonly used workflows for whole genome sequencing including high depth and low coverage sequencing: a miniaturized KAPA HyperPlus workflow (Roche, Li *et al*. 2021), a miniaturized DNA Prep workflow (Illumina, Pillay *et al*. 2023), a miniaturized IDT kit, and a full sized IDT kit. The Illumina DNA Prep kit differs from the others as it is tagmentation-based, as opposed to ligation-based (**Table 1**). Tagmentation can lead to bias depending on the GC content of the sample (Sato *et al*. 2019); however, less bias is observed in the DNA Prep kit than other tagmentation reactions such as Nextera XT (Gunasekera *et al*. 2021). While there are small differences in the proprietary enzymes used for the ligation-based kits and differences in the adapters, the ligation-based kits are very similar in terms of lab protocols. Additionally, Roche uses full length adapters while Illumina and IDT use stubby adapters; whether end repair is allocated a separate step in the protocols is another point of difference between kits (**Table 1**). It should be noted that full length adapters are also available from IDT.

**Table 1:** Key differences between kits.

| Kit | Kit Type | Fragmentation<br>& End Repair | Adapter Used | Size of<br>Reaction |
| --- | --- | --- | --- | --- |
| Roche mini | Ligation | Separate steps | Full length | Miniature |
| Illumina mini | Tagmentation | N/A | Stubby | Miniature |
| IDT | Ligation | One step | Stubby/Full length | Full Sized |
| IDT mini | Ligation | One Step | Stubby/Full length | Miniature |

Miniaturization of library preparation reactions is one approach to reducing costs while maintaining effective library generation (Lai *et al*. 2020; Levine 2019, Mayday *et al*. 2019; Ogiso-Tanaka *et al*. 2023; Pillay *et al*. 2023). Ogiso-Tanaka *et al*. (2023) found that miniaturization reduced 86.8% of reagent usage for amplicon-type libraries. Miniaturization involves reducing reaction sizes, usually with the use of automation to facilitate liquid handling and dispensing, without compromising library yield or quality. Upfront capital expenditure factors into cost decisions when choosing a library preparation workflow, which can be offset by reducing the cost of library preparation, and miniaturization may not be possible with all types of liquid handlers and dispensers, as sub-microliter volumes need to be dispensed and handled, or they may be unavailable and performing a full sized reaction may be necessary. Comparisons of automation strategies for next generation sequencing have been covered in detail elsewhere (Hess 2020). Previously, we reported using a miniaturized version of Roche’s KAPA HyperPlus library preparation kit successfully (Li *et al*. 2021) using this type of automation. The miniaturization of the Illumina DNA Prep method has also been described (Pillay *et al*. 2023). In terms of the actual data produced from sequencing these libraries, total sequencing coverage and duplication rates are particularly important for low coverage sequencing, since low coverage reads are typically imputed against a reference panel to obtain genotype calls. High duplication rates lead to wasted sequencing effort, and low coverage sequencing is especially sensitive to reduced output given its sparse nature. Reduced per-sample coverage (whether due to duplication or other causes) can have a direct impact on imputation performance (Li *et al*., 2021).

In this manuscript, we address the gap in available guidance for low coverage whole genome sequencing by conducting a controlled evaluation of library preparation kits, and assessing them on key metrics including cost, time, coverage, duplication rate, and imputation accuracy. In doing so, we aim to provide data-driven guidance on the choice of preparation kit and to provide a resource for researchers considering low coverage sequencing as an assay.

## METHODS

### Library Preparation and Miniaturization

96 human DNA samples were sourced from the NIGMS Human Genetic Cell Repository at the Coriell Institute for Medical Research (Supplemental Table 1) as high quality sequencing data are available via the 1000 Genomes Project (Auton *et al*. 2015) which is useful for evaluating imputation accuracy. Samples were diluted in low-TE buffer and were subjected to 3 different miniaturized library prep methods: KAPA HyperPlus (Roche, KK8514), DNA Prep (Illumina, 20060059), the library prep kit from IDT, with adaptations for low reaction volumes. Miniaturization involved testing libraries at 1/6th (IDT mini) or 1/8th (Roche mini, Illumina mini) of reaction volume tailoring input DNA to each kit between 24-45 ng total (**Table 2**), and running according to the manufacturer’s instructions, with the exception of fragmentation time for the IDT mini where 4 minutes with 1X reagent K2 was used as over-fragmentation was observed when using the recommended 8 minutes (**Supplementary Figure 1**). The IDT kit was also run according to the manufacturer’s recommendations as full-sized reactions, with the above change to the fragmentation time.

**Table 2:** Input recommendations and number of PCR cycles used for each kit.

| Kit | Input<br>Used | Input Recommendations | Number of PCR<br>Cycles Used |
| --- | --- | --- | --- |
| Roche mini | 44 ng | 22 ng minimum, normalized to 5-10 ng/uL | 6 |
| Illumina mini | 24 ng | 8 ng/μL with a minimum volume of 3μL | 5 |
| IDT | 45 ng | 100 pg–1 μg | 6 |
| IDT mini | 45 ng | 100 pg–1 μg | 6 |

Libraries were cleaned using a 1.0X SPRI bead clean-up (Beckman Coulter, B23319) and quantified via Quant-iT PicoGreen dsDNA Assay (Thermo Fisher Scientific, P7589) on a GloMax Explorer GM3500 (Promega) followed by running a selection of libraries using the dsDNA Quantitation, 1X High Sensitivity Assay (Thermo Fisher Scientific, Q33231) on a Qubit 3.0 (Thermo Fisher Scientific) and inferring the concentration of the remaining libraries. Miniaturized libraries were prepared using automation, the Mantis (Formulatrix) liquid dispenser and the BRAVO (Agilent) liquid handler, while full sized libraries were prepared by hand.

### Library Pooling and Sequencing

Libraries were diluted to the same concentration using Resuspension Buffer (RSB, 0.1mM EDTA, 10mM Tris-HCl) and pooled at equal volumes, one pool per kit. Pools were then size selected by doing a 0.7X left sided SPRI (retaining beads) followed by a 0.56X right sided SPRI (discarding the beads) and then a 1.24X left sided SPRI (retaining beads). Size selected pools were measured by Qubit and Fragment Analyzer (Agilent) to determine concentration and size of the pooled libraries, then sample pools were diluted to 2nM in RSB plus Tween 20 (Illumina, supplied with NextSeq reagents). 2nM pools for each kit were prepared for sequencing on a NextSeq 2000 (Illumina) according to the manufacturer’s instructions and sequenced to ∼0.5X coverage per sample.

### Analysis

Samples from each library preparation kit were mapped to the Human reference genome GRCh38/hg38 (GCA_000001405.15) using bwa mem. Subsequently, the samples were imputed against a high-quality haplotype reference panel known as the HGDP1KG (Koenig et al., 2023), using the GLIMPSE v2 software package (Rubinacci et al., 2023). We provide genotype likelihoods as input to GLIMPSE, calculated from the read pileup directly, at every site in the reference panel assuming a static sequencing error rate of 0.001. The HGDP1KG panel is a combined resource incorporating both the New York Genome Center’s 1KG panel (NYGC1KG) and the Human Genome Diversity Project (HGDP). It features 4,091 samples and covers 76.4 million variants, which include 67.2 million SNPs and 9.2 million indels. It has been previously demonstrated that this panel outperforms an earlier gold standard, the NYGC1KG (Koenig et al., 2023).

To evaluate the performance and consistency of different library preparation kits, we assessed a range of metrics. These metrics were as follows: total bases sequenced (the total number of bases sequenced prior to de-duplicating), raw coverage, effective coverage, duplication rate, and leave-one-out (LOO) imputed genotype concordance. Effective coverage, calculated as -ln(1 - f_covered) with f_covered being the fraction of sites in the imputation panel covered by at least one read, captures the evenness of sequencing reads across sites in the reference panel. The LOO method is a validation procedure that removes the sample being imputed from the reference panel before imputation, to account for potential bias in accuracy estimates. Additionally, we assessed aggregate imputation r^2^ values using the LOO method across varying reference panel minor allele frequency (MAF) bins for each of the five runs. These MAF bins included 0.0001, 0.0005, 0.00075, 0.001, 0.00125, 0.005, 0.01, 0.025, 0.05, 0.1, 0.15, 0.2, 0.25, 0.3, 0.35, 0.4, 0.45, and 0.5. Imputation r^2^ was calculated with the GLIMPSE2_concordance program using a sample of 10 individuals per run (same individuals for each run). Values reported per MAF bin reflect the aggregate imputation r^2^ across the 10 samples.

## RESULTS

The average total sequencing output per sample across runs prior to deduplication ranged from 1.161 Gb to 1.586 Gb. The IDT samples had the lowest average output, and the samples had the highest (**Figure 2; Table 3**). The raw coverage (bases sequenced divided by genome size) varied from 0.497X to 0.520X. This variance also differed by library type. The non-miniaturized IDT library showed the most variation (sd = 0.627 Gb; 95% CI margin of error = 0.126 Gb), than the others, likely due to differences in library quantification and pooling, and not due to a fault in the kit. The miniaturized IDT libraries had slightly lower effective coverage (mean = 0.272 and 0.271, respectively) compared to Illumina mini (mean = 0.364), full sized IDT (mean = 0.340), and Roche mini (mean = 0.356; **Figure 3; Table 3**). Sample effective coverage was lower for miniaturized IDT regardless of raw coverage (**Figure 3**), indicating that the lowered mean effective coverage observed for this run is not driven by a minority of low output samples and that another factor (or factors) is contributing to reduced effective coverage. We hypothesize about these factors below.

**Table 3:** Summary of key performance metrics across the tested kits. Values indicate mean across all samples. Values in parentheses indicate the 95% margin of error, calculated as 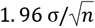.

| Kit | Bases<br>Sequenced | Duplication<br>Rate | LOO<br>Concordance | Raw<br>Coverage | Effective<br>Coverage |
| --- | --- | --- | --- | --- | --- |
| Roche mini | 1.539 Gb<br>(0.071) | 0.105<br>(0.002) | 0.999<br>(5.54e-05) | 0.497X<br>(0.023) | 0.356<br>(0.014) |
| Illumina mini | 1.586 Gb<br>(0.038) | 0.114<br>(0.002) | 0.999<br>(5.12e-05) | 0.512X<br>(0.012) | 0.364<br>(0.001) |
| IDT | 1.161 Gb<br>(0.126) | 0.152<br>(0.003) | 0.999<br>(6.55e-05) | 0.520X<br>(0.041) | 0.340<br>(0.025) |
| IDT mini | 1.570 Gb<br>(0.051) | 0.146<br>(0.002) | 0.999<br>(6.32e-05) | 0.507X<br>(0.017) | 0.271<br>(0.006) |

**Figure 1:**
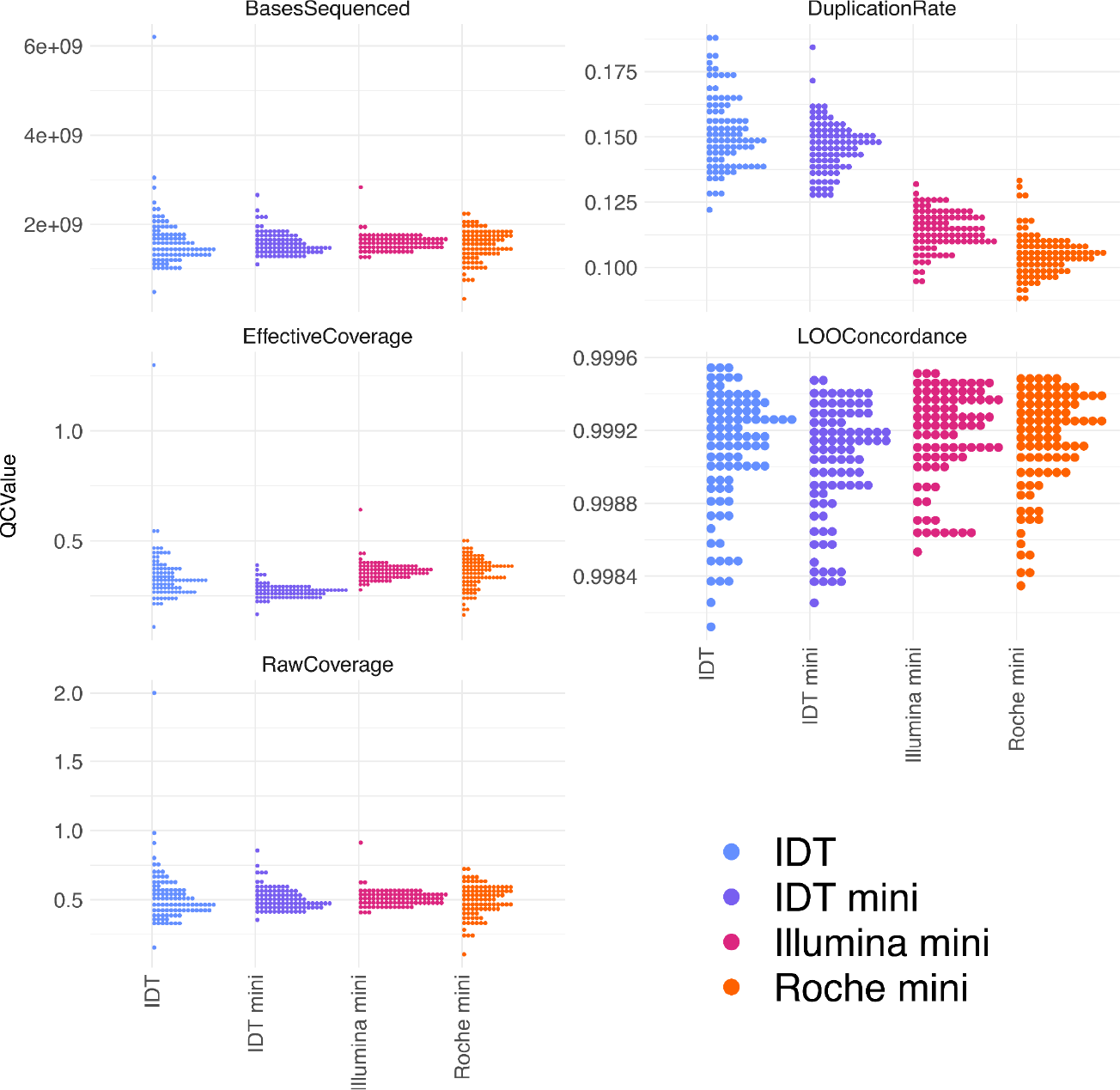
Histograms of performance metrics across tested kits Illumina miniaturized, IDT, IDT miniaturized, and Roche miniaturized. *Note:* y-axes are not shared between panels.

**Figure 2:**
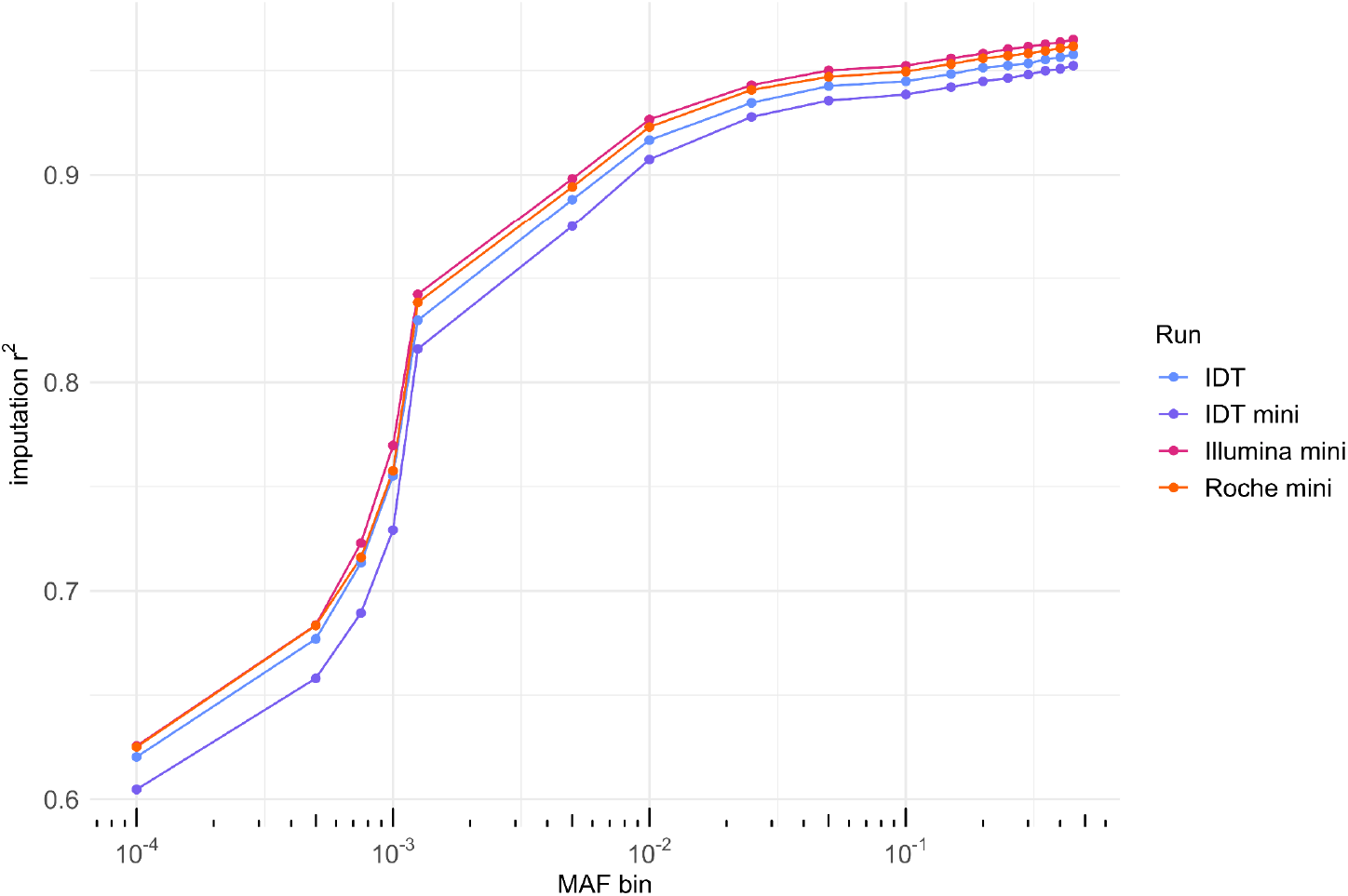
Aggregate imputation r^2^ across minor allele frequency bins for each of the five runs.

**Figure 3:**
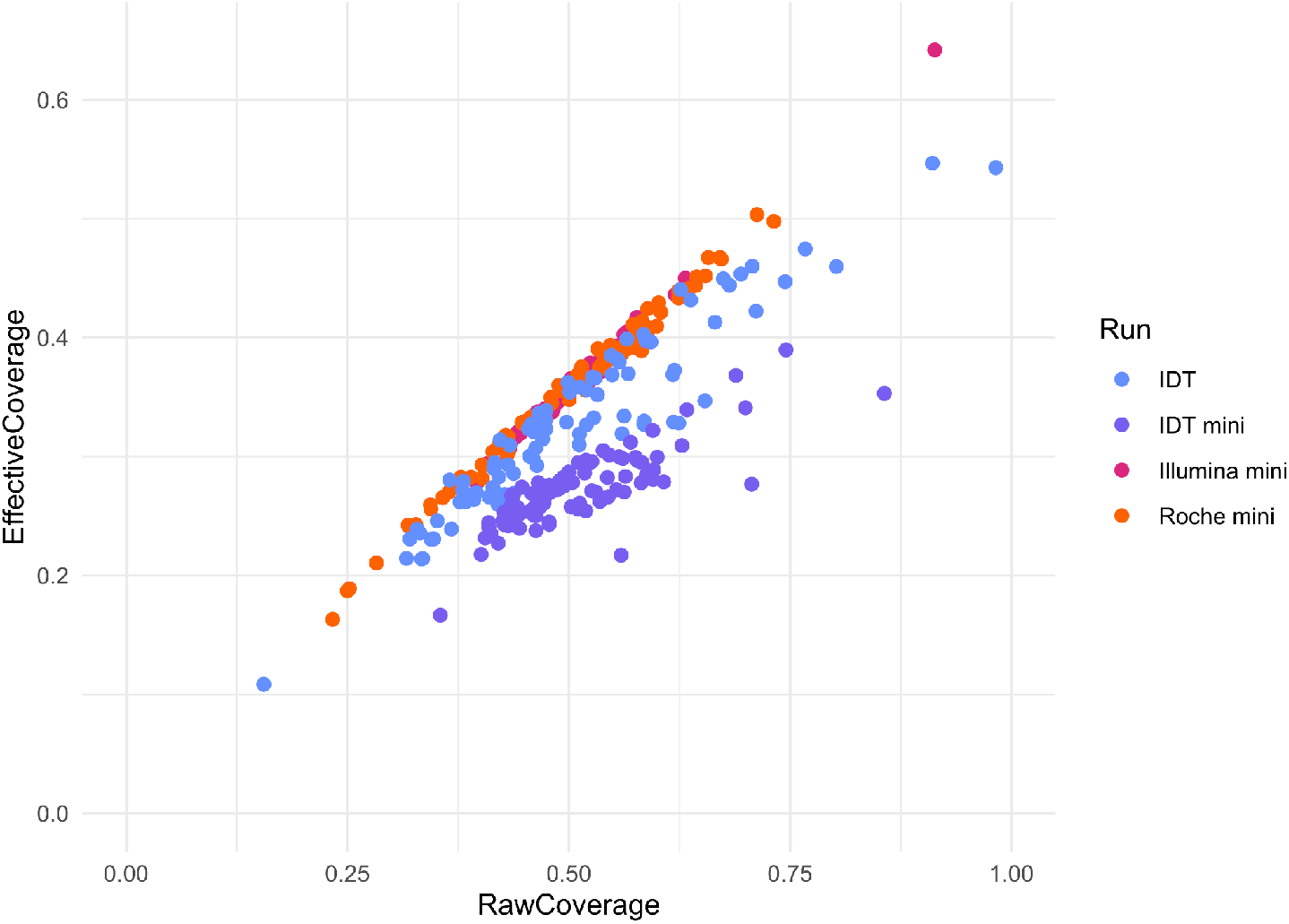
Scatter plot of effective coverage vs raw coverage for each kit indicates that sample effective coverage for IDT mini is lower than than other kits across all values of raw coverage (i.e., that the reduction in effective coverage is not driven by a minority of low output samples).

Duplication rates also varied among the different preparation kits. The IDT protocols, both miniaturized and non-miniaturized, had the highest overall duplication rates, averaging 14.6% and 15.2% respectively. The Roche mini kit showed the lowest duplication rate at 10.5% (**Figure 2; Table 3**). The variance in duplication rates was consistent across libraries, except for the non-miniaturized IDT library, which had slightly higher variability (sd = 0.014; 95% CI margin of error = 0.003; **Figure 2; Table 3**).

In evaluating the impact of library preparation protocol on imputation performance, we found high imputation leave-one-out (LOO) concordance for all libraries (average LOO concordance = 0.99, **Table 3**), reflecting the quality of the HGDP1KG panel used for imputation. The Pearson correlation between imputed and true genotypes, imputation r^2^, was consistent with human imputation standards (Koenig et al., 2023). As expected, the imputation r^2^ increased with minor allele frequency (MAF), with correlations ranging between 0.60 and 0.75 for MAF between 10^-4^ and 10^-3^ to ∼0.96 for MAF above 0.1. Illumina mini samples had, on average, higher r^2^ across MAF bins, although Roche mini was only slightly less performant. IDT mini libraries generally showed the lowest imputation r^2^ for each MAF category and were effectively equivalent. The consensus rank order of performance across MAF bins, from highest to lowest, was: Illumina mini, Roche mini, IDT, and IDT mini.

Miniaturization of the IDT kit resulted in non-statistically significant differences (raw coverage P = 0.55, F = 0.3489, leave one out concordance P = 0.1624, F = 1.9678) in the tested metrics when compared to the full sized IDT kit, with the exception of effective coverage (0.271 vs 0.340, respectively. P = 1.248e-06; F = 25.17) and duplication rate (14.6% vs 15.2%, respectively. P = 8.666e-4; F = 11.458, **Table 3**). Analysis of the insert sizes of these two library types revealed a smaller distribution for IDT mini than IDT (**Figure 4**). 100% of samples successfully generated libraries from the IDT mini kit, with zero repeats needed during library preparation, whereas the other miniaturized libraries needed 4 and 8 samples repeated for Roche mini and Illumina mini, respectively (**Table 4**).

**Table 4:** Summary of lab operations considerations for each kit.

| Kit | Time (Hours) | Cost per Sample* | Agilent BRAVO steps |
| --- | --- | --- | --- |
| Roche mini | 3 | <\$5 | 3 |
| Illumina mini | 2 | <\$5 | 5 |
| IDT | 3 | >\$20 | N/A ^ |
| IDT mini | 3 | <\$5 | 3 |

^Volumes too high for BRAVO automation. \*Not including SPRI, plasticware, tips, etc.

**Figure 4:**
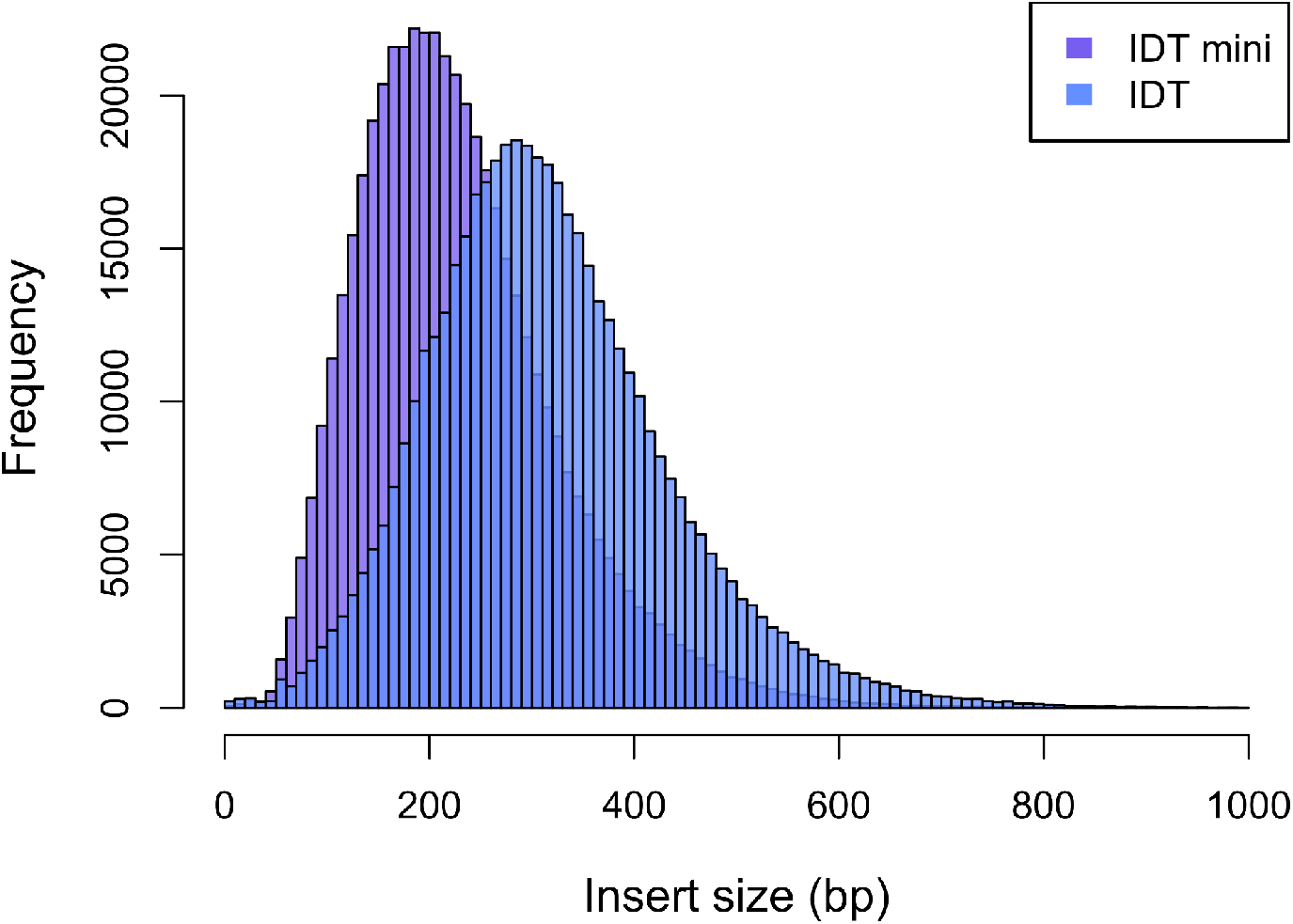
Insert size distribution of IDT and IDT mini for 1M reads of a representative sample (HG00101). Only properly paired reads were used.

Costs per sample for miniaturized libraries are considerably lower than most full sized reactions (**Table 4**). However, this cost saving can be offset by the investment in capital equipment such as liquid handlers. Irrespective of the size of the library, for large sample numbers automation is essential; however investment in liquid handlers for full sized libraries may exceed the pricing for the liquid handlers used in the miniaturized libraries. Miniaturized libraries were <$5 per sample for the kit alone, while the full sized IDT was the most expensive at >$20 per sample (**Table 4**). Miniaturizing IDT brought the cost per sample down to <$5 (**Table 4**).

From a lab handling perspective, all the library kits are easy to prepare and take under 3 hours. The Illumina mini, a tagmentation based assay, is the fastest to prepare and requires the least handling time (2 hours) compared to the ligation based kits (3 hours): Roche mini, IDT, and IDT mini (**Table 4**). Miniaturized libraries can be prepared by hand, but were done using liquid dispensing and liquid handling for this evaluation. 3 steps were needed on the BRAVO for the ligation based kits, however the Illumina mini required 5 steps (**Table 4**). There are trade offs and the decision of which workflow to use will depend on existing equipment and available capital for automation.

## DISCUSSION

Low coverage whole genome sequencing with imputation can be used in place of high-density SNP genotyping arrays and can provide additional insights and power for downstream analyses such as GWAS, PRS, and complex trait mapping (Chat *et al*. 2022; Gilly *et al*. (2019); Homburger *et al*. 2019; Li *et al*. 2021; Wasik *et al*. 2021). One key decision when designing experiments involving whole genome sequencing is in determining the optimal library preparation method: maximizing output with low levels of duplicates, while keeping in mind factors such as time and cost per sample. In particular for GWAS, as sample numbers have now climbed in excess of 1 million (Jansen *et al*. 2019; Lee *et al*. 2019), overall cost is of utmost importance. Here, we described a controlled evaluation of library preparation methods for Illumina low coverage whole genome sequencing.

All of the kits performed well in our hands, with an average of ∼0.3X effective coverage. Differences between the kits in terms of effective coverage may have been due to differences in duplication rates, normalization, or amounts loaded on the sequencer. Differences in variation in the effective coverages may be due to how the quantification of libraries was estimated before pooling. Previous studies report 0.5-1.0X coverage for low coverage sequencing studies for achieving high LOO concordance (Gilly *et al*. 2019; Li *et al*. 2021); however, we previously showed that 0.4X coverage was sufficient for achieving LLO concordance of >0.98. (Wasik *et al*. 2021). The LOO concordance for ∼0.3X coverage in this study shows a 0.99 value across the board (**Table 3**), indicating 0.3X is sufficient for genotyping.

Performance in terms of imputation r^2^ revealed some differences in imputation accuracy between the evaluated kits (**Figure 2**). Namely, these results show that samples prepared with the IDT Mini kit had on average lower correlations to their truth genotypes across all minor allele frequencies. We mainly attribute this drop in performance to the higher duplication rates (**Figure 1**) and decrease in insert sizes (**Figure 4**) observed for this method causing reductions in effective coverage (**Figure 3**). Read duplication results in less sites sequenced across the genome (i.e., lower effective coverage) and thus can have a direct impact on imputation performance. Additionally, the IDT mini fragmentation steps resulted in smaller than expected insert sizes (**Figure 4**), leading to the paired end 150bp reads partially overlapping each other for a substantial fraction of the inserts. This, in theory, contributes to lowered effective coverage, since a fraction of each insert gets sequenced twice (once by read 1 and once by read 2) rather than sequencing effort being distributed across distinct sequences within the insert. For high coverage mutation studies, sequencing across the DNA duplex with both reads is preferable to increase confidence of the call, while in low pass genotyping studies, covering more of the genome improves the power. Optimizing the fragmentation time for the miniaturized reaction to increase fragment size would likely increase effective coverage.

Duplication rates varied between 10.5% (Roche mini) and 15.2% (IDT), with the IDT mini also scoring on the higher end (14.6%, **Table 3**). Rochette *et al* (2023) suggest that PCR duplication rate in sequencing is largely determined by sample quality and complexity (number of sequenceable fragments) and sequencing depth, with PCR cycles playing a secondary role. In our case as the same source DNA was used and sequencing depth is roughly the same for each kit type, the higher duplication rates seen here may be due to the number of cycles used, or may be due to differences in enzyme efficiencies between the kits. As the same number of cycles was also used for the Roche mini (6, **Table 2**) which had lower duplication rates than the IDT kits (**Table 3**), the enzymes and ligation efficiency of the IDT kits may be lower.

Miniaturization of the IDT kit was successful, with the only statistically significant differences being in effective coverage and duplication rate, where the full sized IDT kit performed better in part due to the fragmentation size differences (**Figure 4**). Optimization of the IDT kits could reduce or eliminate these differences. In particular the IDT mini kit needed no repetition of samples where other miniaturized kits required repeat of dropouts (**Table 4**). Costs are substantially reduced when miniaturizing library preparation (Lai *et al*. 2020; Levine *et al*. 2019; Ogiso-Tanaka *et al*. 2023;). We found an 83.3% reduction in usage by miniaturizing the IDT kit (**Table 4**), although most consumables, excluding SPRI beads, would remain the same. This is in line with previous attempts at miniaturization (86.8% reduction in reagent usage, Ogiso-Tanaka *et al*. 2023), and represents significant cost savings.

Finally, from a lab handling perspective, we evaluated the kits on the basis of ease of use. The Illumina mini requires the shortest amount of time, ∼2 hours, as it is tagmentation-based, while the other kits are ligation-based and require ∼3 hours (**Table 4**). The trade off is in the number of steps using automation for each: tagmentation requires washing of the beads and additional steps not required by ligation and thus takes additional steps (5 vs 3, **Table 4**). This makes the protocol more “hands on” as manual set up of the BRAVO and transferring from the Mantis requires physical input, although use of different automation equipment, such as including a liquid dispenser within the liquid handler, could reduce this.

Overall, the performance of all of the kits was impressive: high r^2^ and LOO concordance was observed for all. Therefore, deciding on a library preparation kit to use will depend on several factors, including, but not limited to: the automation instruments already in place, which may dictate the volumes used, the time for preparing libraries and hands on hours, especially if not using automation, whether full length adapters are preferable to your workflow, or the use case. Preparing miniature reactions by hand necessitates pipetting sub-microliter volumes and is not recommended, in this case choosing a full sized kit would be advantageous. For the fastest prep, Illumina mini offers a two hour protocol. For full length adapters, compatible with PCR-free workflows, the Roche mini or the IDT kits are an ideal choice. IDT also offers a deceleration module for fragmentation for generating large fragments for long-read sequencing, which could be useful if sequencing with Nanopore or PacBio.

**Supplementary Figure 1:**
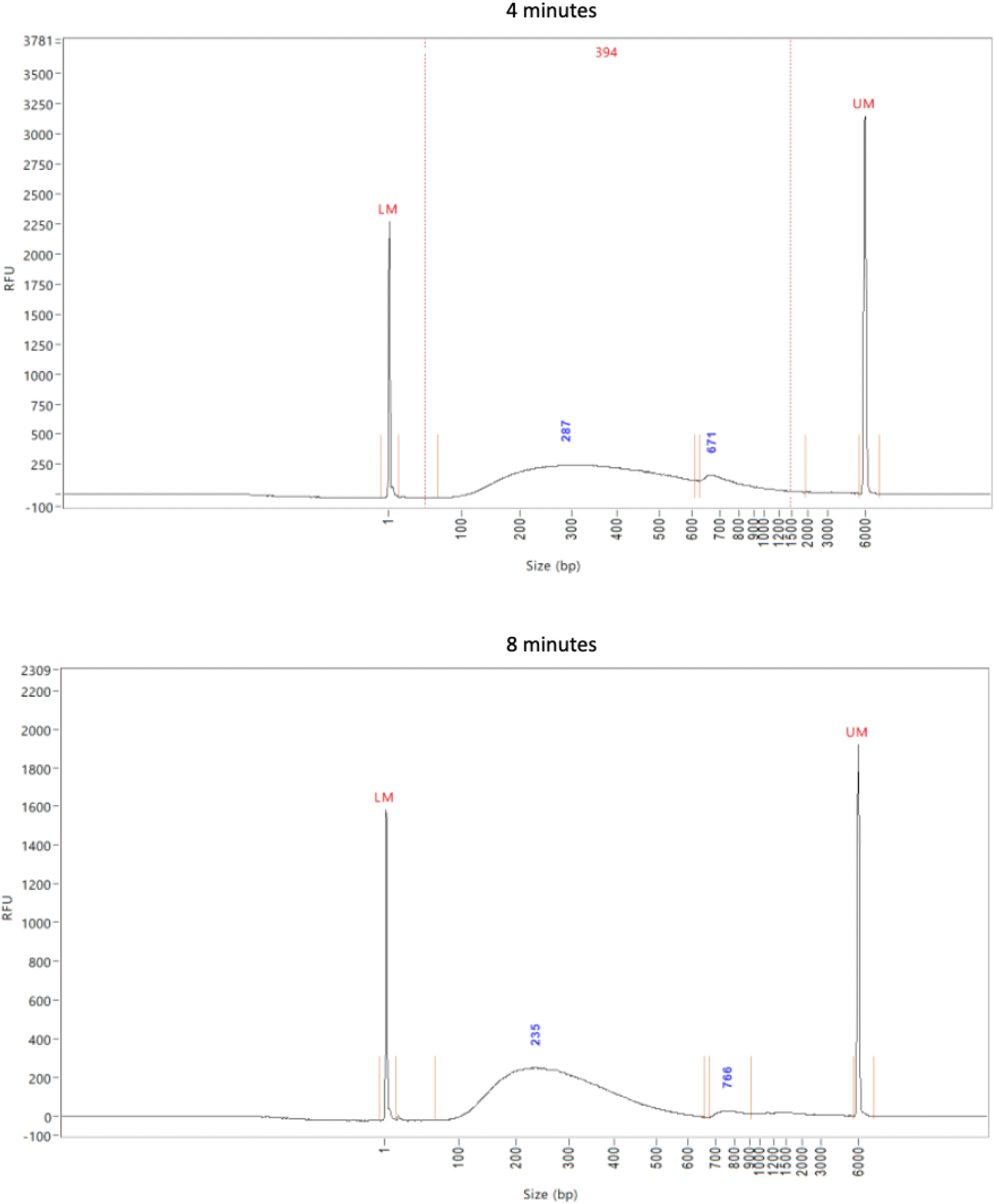
IDT full sized kit, DNA fragment size distribution for different fragmentation times. 2X of K2 solution was used for these samples, 1X was also tested (data not shown).. *LM -Lower Marker, UM - Upper Marker, RFU - Relative Fluorescence Units*.

**Supplementary Table 1:**
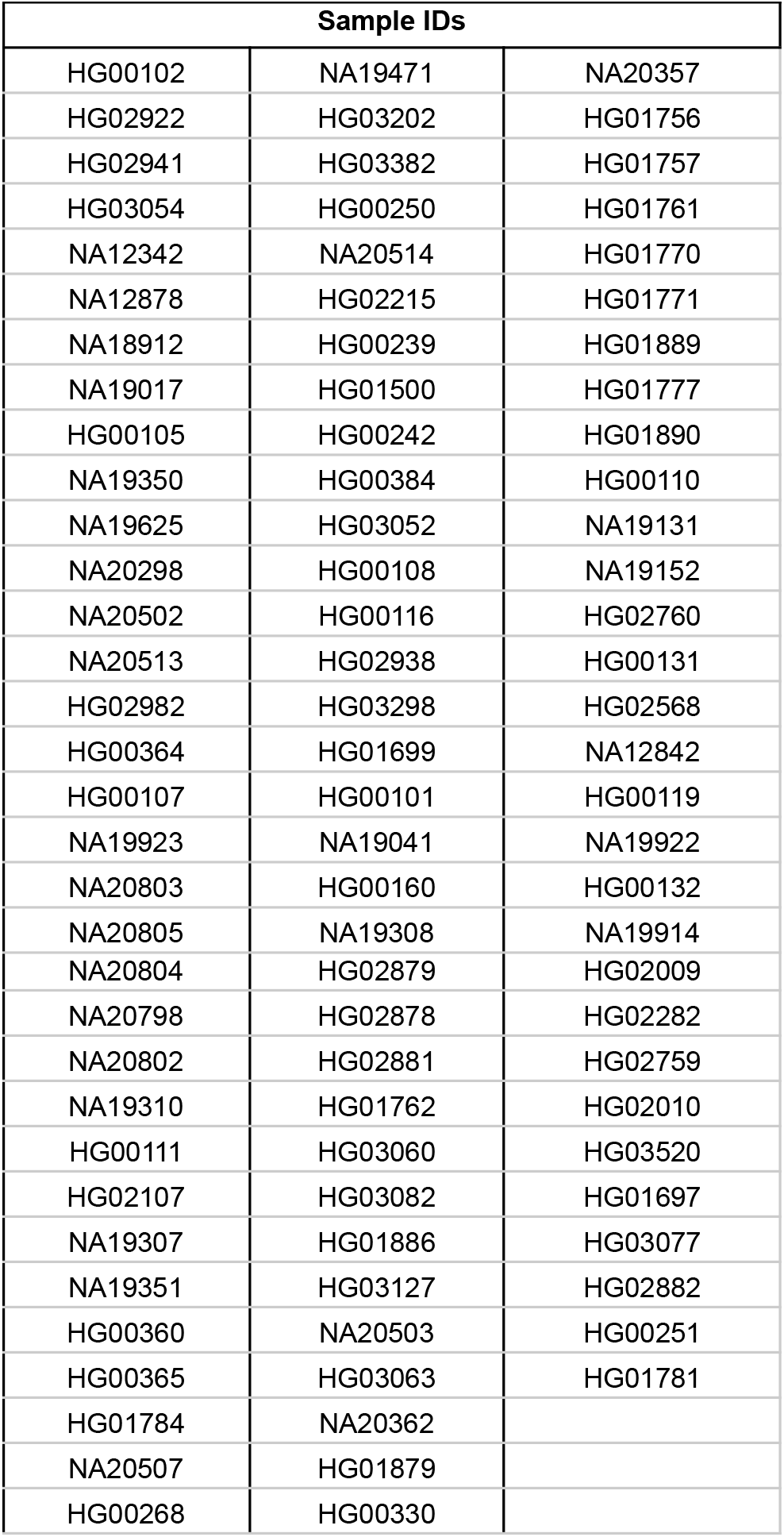
Sample IDs for Coriell samples

## Notes

### Competing Interest Statement

CMS, MJSG, JYP, & JHL were all employees of Gencove, Inc. at the time of writing.

